# EFR3 and phosphatidylinositol 4-kinase IIIα regulate insulin-stimulated glucose transport and GLUT4 dispersal in 3T3-L1 adipocytes

**DOI:** 10.1101/2021.06.02.446733

**Authors:** Anna M. Koester, Kamilla M. Laidlaw, Silke Morris, Marie F.A. Cutiongco, Laura Stirrat, Nikolaj Gadegaard, Eckhard Boles, Hannah L. Black, Nia J. Bryant, Gwyn W. Gould

**Author notes:** Equal contribution (Anna M. Koester, Kamilla M. Laidlaw). Correspondence to joint communicating authors or.

## Abstract

Insulin stimulates glucose transport in muscle and adipocytes. This is achieved by regulated delivery of intracellular glucose transporter (GLUT4)-containing vesicles to the plasma membrane where they dock and fuse, resulting in increased cell surface GLUT4 levels. Recent work identified a potential further regulatory step, in which insulin increases the dispersal of GLUT4 in the plasma membrane away from the sites of vesicle fusion. EFR3 is a scaffold protein that facilitates localisation of phosphatidylinositol 4-kinase type IIIα to the cell surface. Here we show that knockdown of EFR3 or phosphatidylinositol 4-kinase type IIIα impairs insulin-stimulated glucose transport in adipocytes. Using direct stochastic reconstruction microscopy, we also show that EFR3 knockdown impairs insulin stimulated GLUT4 dispersal in the plasma membrane. We propose that EFR3 plays a previously unidentified role in controlling insulin-stimulated glucose transport by facilitating dispersal of GLUT4 within the plasma membrane.

## Introduction

Insulin stimulates glucose transport in adipocytes and muscle. This is mediated by the regulated delivery of the glucose transporter GLUT4 from intracellular stores to the plasma membrane (PM). A wealth of information has underpinned our understanding of intracellular GLUT4 trafficking and the mechanism by which GLUT4-containing vesicles fuse with the PM (Gould et al., 2020; Jaldin-Fincati et al., 2017; Klip et al., 2019; Saltiel, 2021; Santoro et al., 2021).

More recently, it has become clear that organisation of GLUT4 within the PM is also subject to regulation by insulin; GLUT4 is found here in both relatively stationary clusters and as freely diffusible monomers (Stenkula et al., 2010). In the absence of insulin, GLUT4 is retained in clusters at the site of fusion; these clusters nucleate clathrin assembly and are thought to represent sites of GLUT4 internalisation (Stenkula et al., 2010). Insulin stimulation was observed to mediate increased delivery of GLUT4 to the PM accompanied by increased dispersal of GLUT4. Two distinct types of GLUT4-vesicle exocytosis were identified: ‘fusion-with-release’ events dispersed GLUT4 within the PM whereas ‘fusion-with-retention’ events retained GLUT4 molecules at the fusion site. In the basal state ~95% of events observed were ‘fusion-with-retention’ and insulin led to a 60-fold increase in ‘fusion-with-release’ events in 2-3 minutes (Stenkula et al., 2010). A follow-up study investigated GLUT4 cluster retention and the molecular dynamics guiding GLUT4 exchange in the PM, and insulin’s effects on GLUT4 organization at high resolution (Lizunov et al., 2013a). Isolated adipocytes were transfected with photo switchable HA-GLUT4-EOS and live cell single molecule tracking was performed with fluorescence photoactivation localization microscopy. The data suggest that insulin had three separable effects that contribute to the shift of GLUT4 molecules within the PM from a clustered to a more dispersed state. Firstly insulin shifted the fraction of dispersed GLUT4 upon delivery to the PM, secondly insulin increased dissociation of GLUT4 monomers from clusters, and lastly insulin decreased the rate of GLUT4 endocytosis (Lizunov et al., 2013a).

The concept of insulin-stimulated GLUT4 dispersal has been further supported using other imaging methods (Gao et al., 2017; Lizunov et al., 2013a). These approaches confirmed a shift in the distribution of the spatial organisation of GLUT4 towards a more dispersed state following insulin stimulation. Interestingly, insulin resistance was found to induce a more clustered distribution of GLUT4, with more molecules per cluster, implying that the regulation of post-fusion GLUT4 distribution may be impaired in disease (Gao et al., 2017). Such studies underscore a need to identify mechanisms that regulate GLUT4 spatial distribution. Here we report the identification of one such potential mechanism.

Here we identify EFR3 as a key regulator of GLUT4 dispersal in the PM. EFR3 is a palmitoylated protein responsible for PM localisation of phosphatidylinositol 4-kinase type IIIα (PI4K-IIIα) (Baird et al., 2008; Nakatsu et al., 2012a; Vijayakrishnan et al., 2009; Wu et al., 2014). We show that both EFR3 and PI4K-IIIα are required for insulin-stimulated glucose transport. Using direct stochastic reconstruction microscopy (dSTORM) we demonstrate that knockdown of EFR3 reduces insulin-mediated dispersal of GLUT4. Furthermore, we find that levels of both EFR3 and PI4K-IIIα are elevated in rodent models of insulin resistance and present a model in which EFR3 serves as a regulatory hub controlling insulin-stimulated GLUT4 dispersal in the PM.

## Results

### EFR3 is implicated in GLUT4 trafficking

When heterologously expressed in the yeast *S. cerevisiae* the mammalian glucose transporter GLUT4 is retained intracellularly, underscoring the conservation of molecular machinery required for regulated trafficking (Shewan et al., 2013). Even when localised to the cell surface, GLUT4 does not efficiently transport glucose suggesting that regulation of GLUT4 within the PM might be important (Kasahara and Kasahara, 1997; Wieczorke et al., 2003). We carried out a genetic screen to select mutations in the yeast genome that enable mammalian glucose transporters to support uptake of glucose into yeast cells lacking their own endogenous hexose transporters (Wieczorke et al., 2003). Expression of GLUT4 was unable to support growth on glucose of yeast lacking endogenous hexose transporters unless they also carry the recessive mutant *fgy1-1* allele (Wieczorke et al., 2003). *fgy1-1* is a mutant allele of *EFR3* (Wieczorke and Boles, personal communication; (Schmidl et al., 2020).

*EFR3* is required for Stt4-containing phosphoinositide kinase patch assembly at the PM (Baird et al., 2008). Stt4 is the yeast orthologue of the human type III phosphatidylinositol-4-kinase IIIα which localises to punctate dots on the PM (Nakatsu et al., 2012a). Efr3 is similarly found at the cell surface (Nakatsu et al., 2012a). This localisation requires the addition of palmitoyl moieties to one or more cysteine residues towards the N-terminus. Intriguingly, the *D. melanogaster* Efr3 orthologue Rolling-Blackout is required for synaptic vesicle exocytosis (Vijayakrishnan et al., 2009), although its precise role is yet to be defined.

These data suggested to us that EFR3 and PI4K-IIIα may play a role in the regulation of glucose transport in response to insulin and prompted us to interrogate the role of EFR3 in insulin regulated GLUT4 trafficking.

### EFR3A is the major isoform expressed in adipocytes

There are two EFR3 paralogs in higher eukaryotes, EFR3a and EFR3b. We used RT-PCR to ascertain which are expressed in our experimental model 3T3-L1 cells. Figure 1A shows that EFR3a is highly expressed in 3T3-L1 fibroblasts and adipocytes; levels of EFR3b are 30-100-fold lower. EFR3 functions as a part of a protein complex that delivers active PI4K-IIIα to the PM; EFR3 associates with the PM via palmitoylation (Baird et al., 2008; Nakatsu et al., 2012a). We therefore asked whether EFR3 and PI4K-IIIα were PM localised in 3T3-L1 adipocytes, and whether insulin modulated this distribution. Subcellular fractionation revealed that, as in yeast (Bojjireddy et al., 2015; Nakatsu et al., 2012a), both EFR3 and PI4K-IIIα are predominantly found in PM-enriched fractions with some within intracellular fractions. In contrast to GLUT4, EFR3 and PI4K-IIIα do not exhibit altered fractionation profiles in response to insulin (Figure 1B). The antibodies used to detect EFR3 cannot distinguish between EFR3a and b but given the RT-PCR data in Figure 1A this is likely to represent EFR3A. Transient expression of mCherry-tagged EFR3a in 3T3-L1 adipocytes also revealed predominantly PM localisation, consistent with EFR3 localisation in other cell types (Baird et al., 2008; Nakatsu et al., 2012b; Wu et al., 2014). In agreement with the fractionation data, no significant alteration of this pattern was evident in insulin-stimulated cells (Figure 1C).

**Figure 1.**
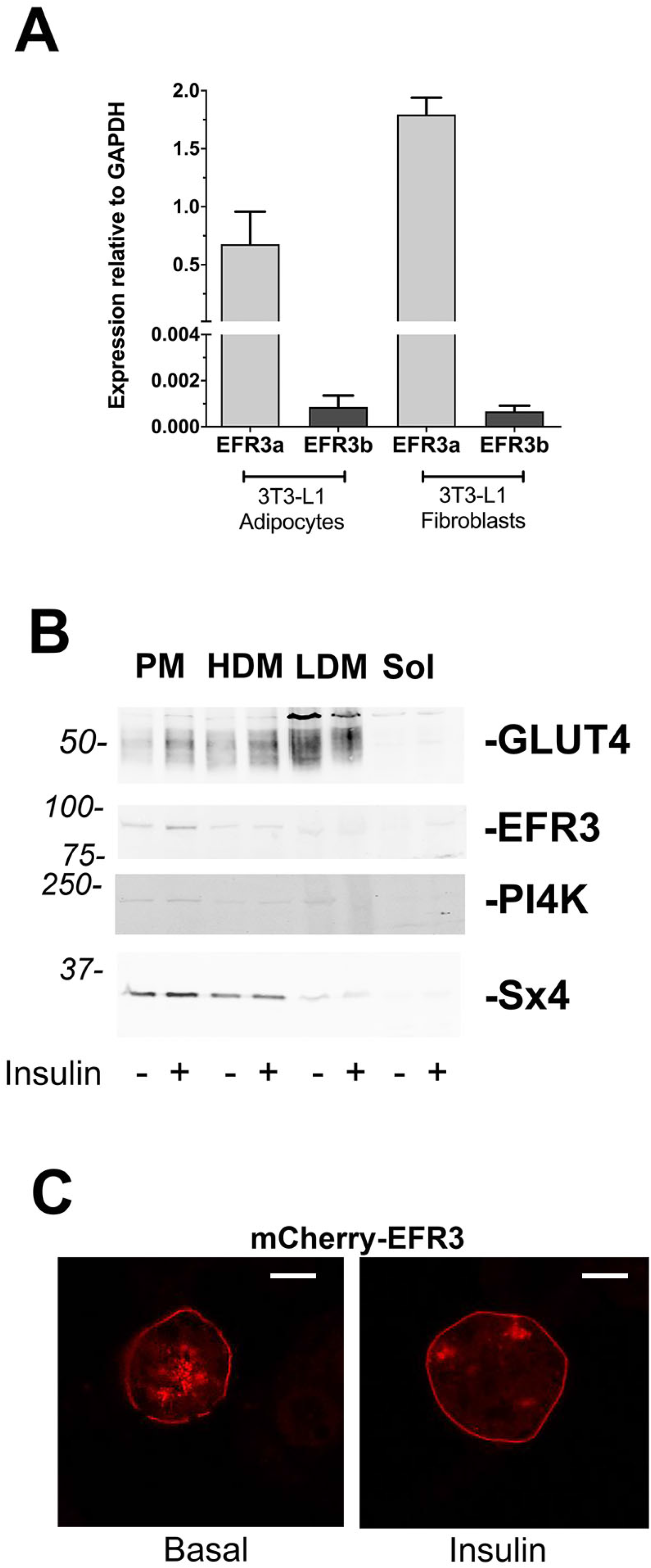
EFR3a is the major isoform in 3T3-L1 adipocytes and is localised to the plasma membrane. **Panel A** shows RT-PCR analysis of the expression of EFR3a and EFR3b normalised to GAPDH in 3T3-L1 fibroblasts and adipocytes. Data presented is from triplicate biological repeats each with at least three technical replicates shown (mean and S.D.) EFR3a is the predominant isoform in 3T3-L1 fibroblasts and adipocytes. **Panel B** shows a subcellular analysis of 3T3-L1 adipocytes treated with or without 100 nM insulin for 20 minutes and separated into PM (PM)-enriched, high density microsomes (HDM), low density microsomes (LDM) and soluble protein (Sol) fraction, which were then immunoblotted for the proteins indicated; figures at left are approximate positions of MW markers in kDa. In each case, 30 μg of protein was loaded in each fraction. GLUT4 levels increase in the PM fraction and decrease in the LDM in response to insulin. EFR3 and PI4K-IIIα are predominantly PM localised and do not appear to redistribute in response to insulin. Data from a representative experiment is shown, replicated three times with qualitatively similar results. **Panel C** shows the distribution of mCherry-EFR3a in 3T3-L1 adipocytes in the presence and absence of insulin. Scale bar 5 μm, data representative of >30 cells in three experiments.

### EFR3 and PI4K are required for insulin-stimulated glucose transport in adipocytes

To directly assay effects on insulin-stimulated glucose transport we used siRNA to knockdown EFR3 or PI4K-IIIα in 3T3-L1 adipocytes. Immunoblot analysis revealed a reduction in EFR3 levels of 48.2% and PI4K-IIIα of 41.5% in cells electroporated with the siRNA against EFR3 and PI4K-IIIα respectively, compared to scr-siRNA treated cells (Figure 2A). Insulin-stimulated a 11.3 + 3-fold increase in 2-deoxy-D-glucose uptake in cells electroporated with scr siRNA (Figure 2B). Knockdown of EFR3 or PI4K-IIIα reduced this to 2.7 + 0.8-fold and 2.6 + 1.5-fold, respectively (Figure 2B). Although there are many potential explanations for an observed reduction in insulin-stimulated glucose transport, including defective insulin signalling, the localisation of EFR3 to the PM prompted us to test the hypothesis that EFR3 knockdown impairs insulin-stimulated glucose transport by an effect on GLUT4 dispersal.

**Figure 2.**
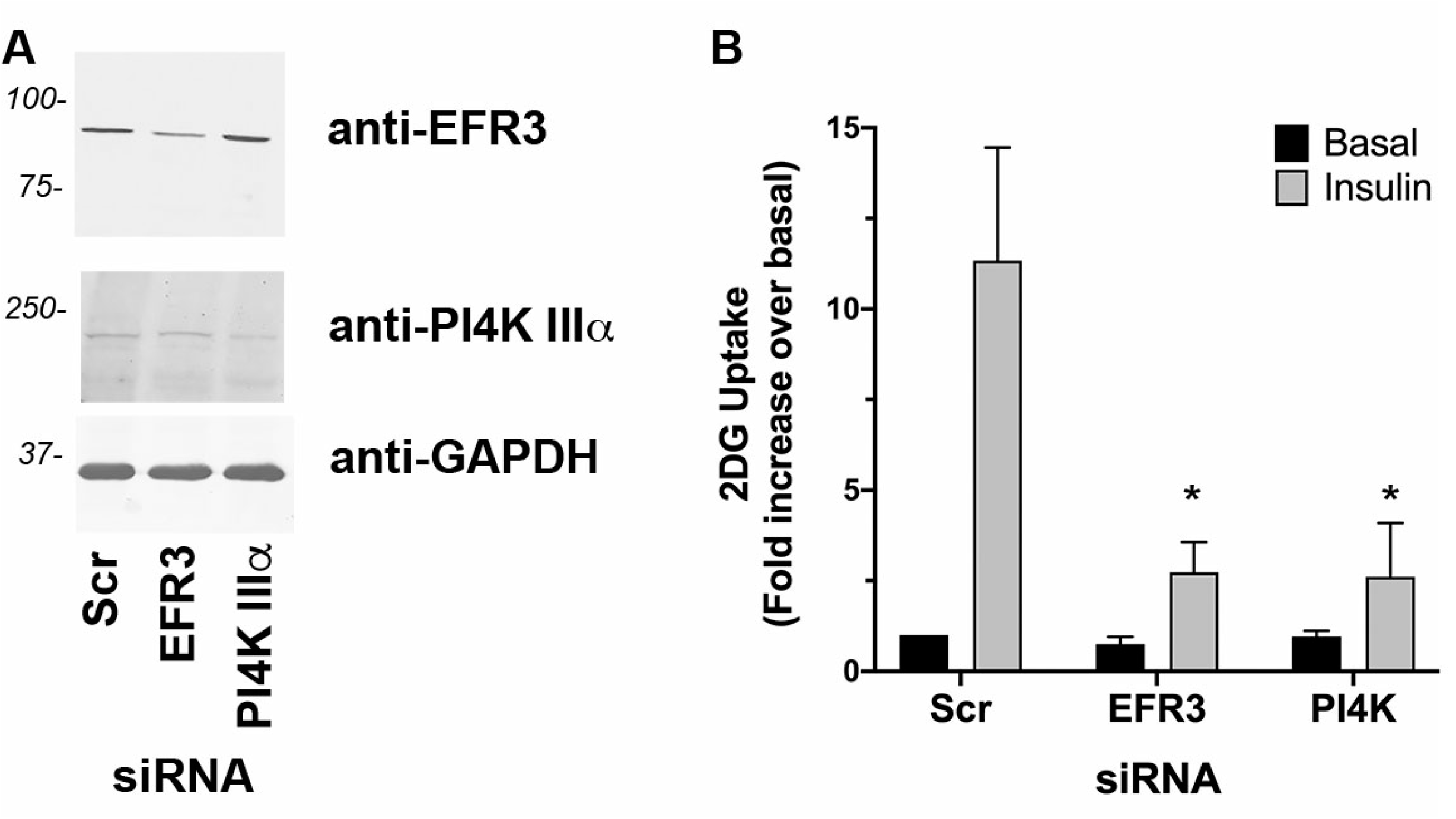
Knockdown of EFR3 impairs insulin-stimulated 2DG transport. **Panel A** 3T3-L1 adipocytes were electroporated with siRNA designed to knockdown EFR3a or PI4K-IIIα, or scrambled control siRNA (Scr) as described. Cells were incubated in serum-free media for 2h, with or without 100 nM insulin for the last 30 minutes then 2DG uptake measured. Because cell numbers varied considerably between experiments, the data is normalised to the rate obtained in the Scr-cells in the absence of insulin. Data shown as mean and SEM of data from at least 3 biological repeats, each with 5 technical replicates. Knockdown of either EFR3 or PI4K-IIIα impaired insulin-stimulated 2DG uptake (*p<0.001 for both). Differences in basal transport rate were not significant. **Panel B** shows a representative immunoblot in which levels of EFR3 and PI4K-IIIα, together with GAPDH, are shown in lysates of cells treated with the indicated siRNA. Data from a representative experiment is shown.

### EFR3 knockdown impairs insulin stimulated GLUT4 dispersal in the PM

Super-resolution imaging approaches have revealed reorganization of GLUT4 in the PM at the single molecule level (Gao et al., 2017; Lizunov et al., 2013a; Stenkula et al., 2010). GLUT4 distribution in the PM was non-homogenous: GLUT4 molecules form clusters or monomers in 3T3-L1 adipocytes. Ripley’s K function spatial analysis revealed that GLUT4 molecules were less clustered after insulin stimulation (Gao et al., 2017). We used a similar dSTORM approach to assess the effects of EFR3 knockdown on GLUT4 distribution in the presence and absence of insulin on samples processed using a standard immunofluorescence protocol with a commercially obtained HA-antibody. While this approach generates consistent imaging results the application of fluorescently labelled antibodies decreases the quantitative accuracy of localization of molecules due to their size, a phenomenon known as antibody linkage error. To control for this, we performed comprehensive spatial point pattern analysis rather than quantifying individual cluster descriptors.

HA-GLUT4-GFP expressing 3T3-L1 adipocytes were electroporated with scr-siRNA or siRNA targeting EFR3, incubated with or without 100 nM insulin for 20 minutes and a dSTORM dataset was collected from cells stained with anti-HA in the absence of detergent, so that only cell surface GLUT4 was immuno-labelled. ThunderSTORM was used to localize each recorded emission of the individual GLUT4 single molecules and generate output files that contain the complete x/y coordinates of all molecule localizations from the numbers of cells reported in the figure legends (Ovesný et al., 2014).

We first qualitatively analysed the data using Hierarchical Density-Based Spatial Clustering of Applications with Noise (Campello et al., 2013; Ester et al., 1996) to identify clusters of GLUT4 in dSTORM images (Figure 3A). In basal cells *HDBSCAN* identified both GLUT4 clusters (coloured structures; Figure 3A) and dispersed GLUT4 molecules (grey dots; Figure 3A). After insulin-stimulation more GLUT4 clusters are visible in the PM and the density of dispersed GLUT4 molecules is increased (Figure 3A). This result is in line with the current literature stating that about 50% of GLUT4 molecules are clustered in the basal state and insulin stimulation induces an increase in the number of GLUT4 clusters in the PM (Stenkula et al., 2010). Knockdown of EFR3 had little effect on the numbers of GLUT4 clusters, but insulin-stimulation did not increase the density of GLUT4 monomers, suggesting that GLUT4 dispersal is impaired.

**Figure 3.**
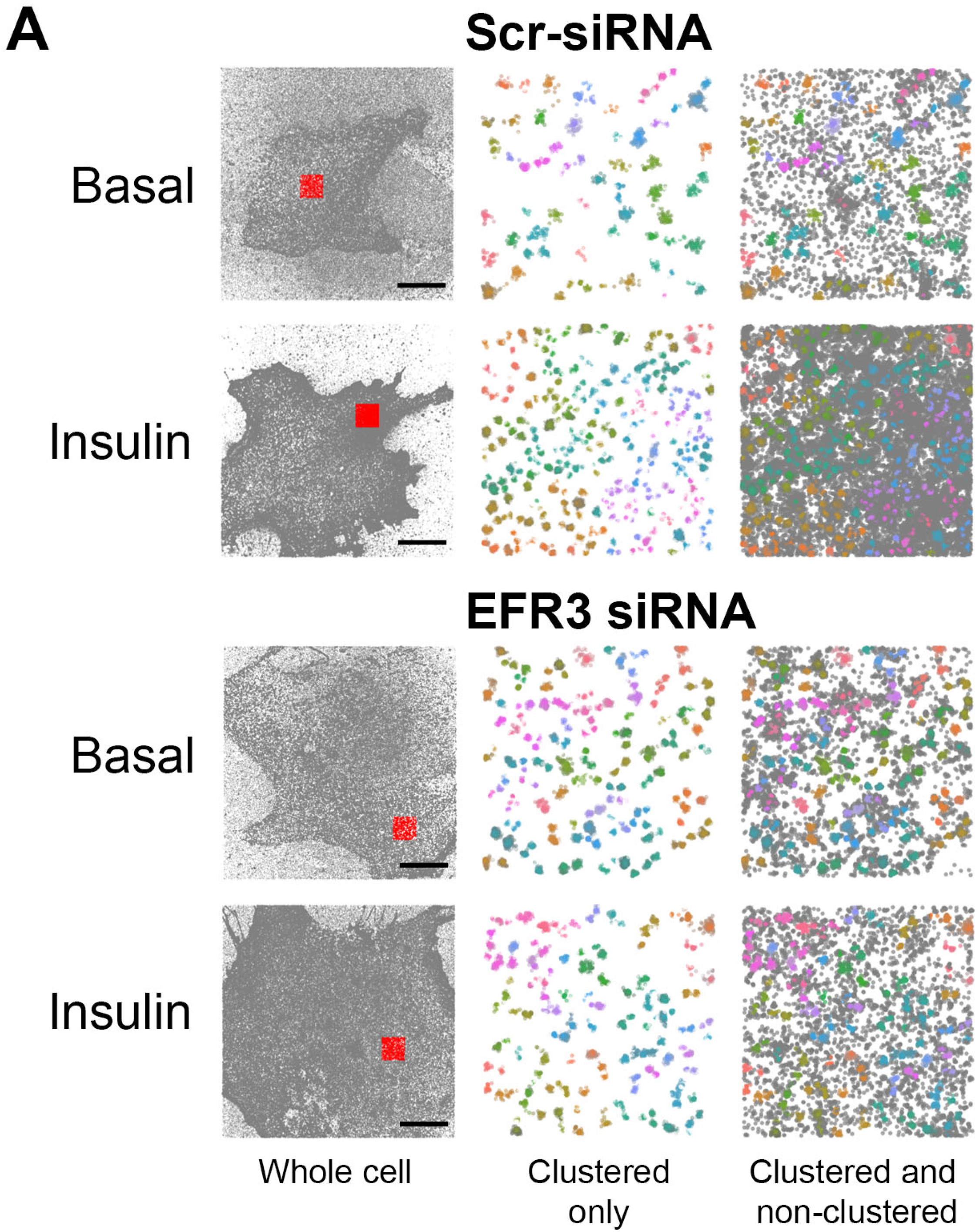
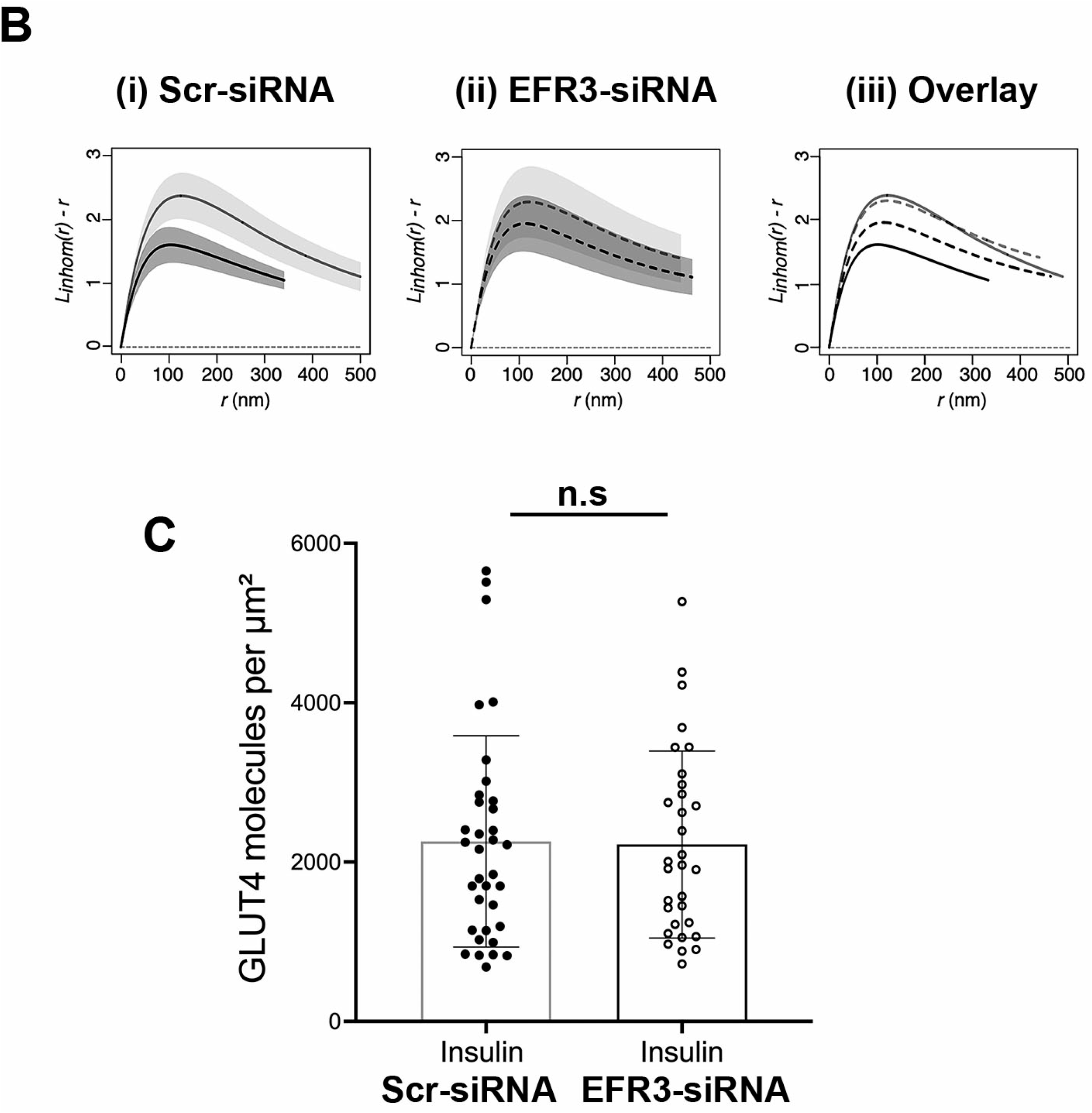
Insulin-stimulated GLUT4 dispersal is impaired upon EFR3 knockdown. 3T3-L1 adipocytes stably expressing HA-GLUT4-GFP were electroporated with siRNA to knockdown EFR3, or a corresponding Scr-siRNA as described. Cells were stimulated with 100 nM insulin for 20 min, or left untreated (Basal), before fixation and staining for surface HA as decribed in *Methods*. dSTORM images were acquired and reconstructions calculated using ThunderSTORM. ***Panel A. Hierarchical density-based spatial clustering of applications with noise analysis for representative control and EFR3 knock-down basal and insulin-stimulated 3T3-L1 adipocytes***. GLUT4 molecule coordinates were processed using a python *HDBSCAN* script written in house. Gray dots indicate single molecule localizations for representative whole cells (left panels). Red boxes indicate region of interests shown in the middle and right panels. The middle panels show coloured molecule clusters identified by *HDBSCAN* within an ROI (‘clustered only’) and coloured clusters plus gray non-clustered molecules identified by *HDSCAN* within an ROI (‘clustered and non-clustered’, right panels) are shown. The representative 4 μm x 4 μm ROI is highlighted by the red square. Scale bars = 10 μm. ***Panel B. L function analysis of GLUT4 molecule clustering in EFR3 knock down basal and insulin-stimulated 3T3-L1 adipocytes***. GLUT4 molecule coordinates were obtained using ThunderSTORM and its spatial pattern was analysed. We analysed the correlation of points using the variance stabilised L function, which is the transformed version of Ripley’s K function. The L function describes how many points (given by L(r)) can be found within a distance r of any arbitrary point. Empirical estimates of the centered L(r) function for (A) control basal and insulin-stimulated cells, (B) EFR3 knock down basal and insulin-stimulated cells, (C) all experimental groups combined. The experiment was carried out independently 4 times on n=10 cells per group. Empirical estimates of the L function from each were pooled together to provide a weighted mean and the 83% (or 1.37σ) confidence interval for each group – see text for details. Complete spatial randomness, modelled from a Poisson process, is indicated by the dashed line. ***Panel C. GLUT4 surface density upon EFR3 knockdown***. HA (surface GLUT4) localisation density was determined from all cells analysed in the presence of insulin. The GLUT4 localisation values do not differ between Scr- and EFR3-siRNA treated groups (n.s.).

To verify this, we used statistical analysis of spatial point pattern data in R (*spatstat*) to compute an estimate for the edge-corrected inhomogeneous Ripley’s L-function. We first confirmed using null hypothesis testing that all the GLUT4 spatial points were significantly different from complete spatial randomness (indicated by the dashed line in Figure 3B). As shown in Figure 3B, the inhomogeneous L(r) peaks at higher values for basal cells indicating more GLUT4 is found clustered at closer distances compared to insulin-stimulated cells, and closely resembles that reported by others (Gao et al., 2017). Knockdown of EFR3 was found to have little effect on the basal distribution of GLUT4, but significantly reduced the dispersal observed in response to insulin (Figure 3B).

We further quantified HA (surface GLUT4) levels from ThunderSTORM datasets (van de Linde, 2019). Strikingly, Figure 3C indicates that the level of GLUT4 at the surface in the presence of insulin is not changed by EFR3 knockdown.

These data are consistent with the idea that depletion of EFR3 has impaired the transition from clustered to dispersed GLUT4 in the PM, independent of any effects on GLUT4 levels.

### EFR3 and PI4K-IIIα levels are increased in cardiac tissue of high fat fed mice

Rodent models of insulin resistance are useful predictors of human disease. We therefore ascertained whether levels of EFR3 and/or PI4K-IIIα are changed in muscle of high-fat fed mice compared to controls using tissue from mice previously characterised and shown to exhibit impaired glucose tolerance and increased body weight (Almabrouk et al., 2018). EFR3 and PI4K-IIIα levels are significantly elevated in cardiac tissue in the high-fat fed group (2- and 1.6-fold, respectively; Figure 4).

**Figure 4.**
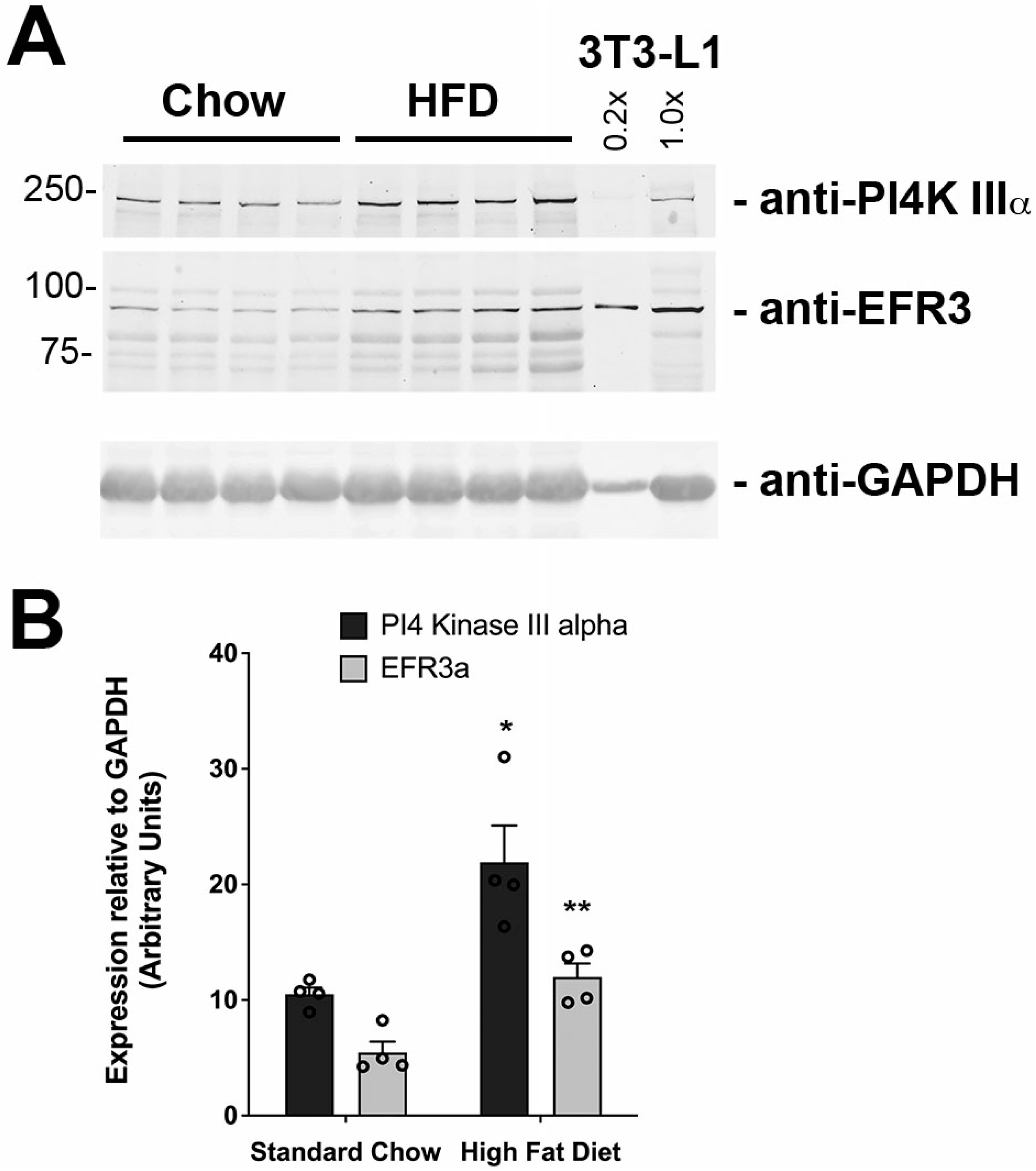
EFR3 and PI4K-IIIα levels increase in rodent models of insulin resistance. Mice fed either standard chow or high-fat diet were as previously characterised (Almabrouk et al., 2018; Mancini et al., 2017). Extracts of samples of cardiac tissue lysates were immunoblotted for EFR3, PI4K-IIIα or GAPDH as indicated. 3T3-L1 adipocyte lysate was used to normalise between blots. Representative immunoblots are shown in **panel A** with the mean data from 4 independent immunoblot analyses presented in **panel B**. * indicates a significant increase in PI4K-IIIα (p=0.012) and ** in EFR3 (p=0.017) in high fat-fed animals compared to controls.

## Discussion

While many of the details regarding the intracellular trafficking of GLUT4 are becoming clearer, much less known about the behaviour of GLUT4 once at the PM. Several lines of investigation suggest that controlling GLUT4 at the PM could be a key facet of insulin action. Koumanov et al developed a cell-free system to study intracellular GLUT4 vesicle fusion with vesicles prepared from the PM of adipocytes; this fusion was cytosol and energy-dependent. Remarkably GLUT4 vesicle fusion with PM-derived vesicles was enhanced 8-fold when PMs were prepared from insulin-stimulated cells, even if internal GLUT4 vesicles and the cytosol were from unstimulated (basal) cells (Koumanov et al., 2005). This study provides compelling evidence for an insulin-regulated event at the PM.

Recent work has suggested that GLUT4 clustering may be an important control mechanism. GLUT4 is thought to function as a monomer in the PM, distinct from other transporters such as GLUT1 which are proposed to form oligomers to regulate functional activity (Hebert and Carruthers, 1992; Zottola et al., 1995). Many studies have argued that GLUT4 is non-randomly distributed within the PM of adipocytes (both primary and 3T3-L1 adipocytes) and muscle, with studies describing a punctate distribution at the cell surface (Karlsson et al., 2002; Lauritzen et al., 2008; Li et al., 2004; Lizunov et al., 2005; Malide and Cushman, 1997). Some attribute this to clustering with clathrin or caveolae, but recent studies argue against this (Stenkula et al., 2010). These studies have recently been extended by imaging approaches which describe GLUT4 clusters in the PM and the ability of insulin to promote GLUT4 dispersal from these clusters (Gao et al., 2017; Stenkula et al., 2010). The observation that insulin-stimulated GLUT4 dispersal is reduced in insulin resistant models (Gao et al., 2017) and the behaviour of GLUT4 vesicles adjacent to the PM is modulated in insulin resistance further emphasises the potential importance of PM-associated GLUT4 events (Lizunov et al., 2013b).

### EFR3 regulates insulin-stimulated glucose transport

Our identification of the scaffold protein EFR3 as a GLUT4 regulatory protein has provided new insight into GLUT4 regulation at the PM. Here we show that EFR3 is predominantly PM localised and knockdown impairs insulin-stimulated glucose transport in 3T3-L1 adipocytes. Our observation that knockdown of the EFR3-associated protein PI4K-IIIα similarly impaired insulin-stimulated glucose transport provides compelling support for a role for EFR3/PI4K-IIIα in the regulation of GLUT4.

### EFR3 regulates GLUT4 dispersal

Data presented here offer the hypothesis that EFR3 may act as a regulator of GLUT4 dispersal. Using dSTORM we examined the distribution of GLUT4 clusters in the PM and used spatial point pattern data in R to compute an estimate for the edge-corrected inhomogeneous Ripley’s L-function. These data reveal that EFR3 knockdown significantly impairs the ability of insulin to promote dispersal of GLUT4 in the PM (Figure 3B). In this analysis, we used anti-HA to label cell surface GLUT4 followed by detection with a labelled secondary antibody. At high resolution, antibodies can significantly limit how well the image reflects the actual structure by increasing the apparent size of visualised structures; for this reason, we have not attempted to quantify the number of molecules per cluster or cluster density. However a visual representation of the data generated using Hierarchical Density-Based Spatial Clustering of Applications with Noise (Figure 3A) is consistent with insulin driving an increase in the number of clusters and an increase in the density of GLUT4 monomers, similar to that reported by others using this approach (Gao et al., 2017). Strikingly, this analysis also clearly reveals that GLUT4 dispersal is impaired upon EFR3 knockdown (Figure 3A,B). To our knowledge, this is the first clue to the mechanism of insulin stimulated GLUT4 dispersal. Quantification of the localisation density of GLUT4 in the PM revealed that the ability of insulin to stimulate GLUT4 translocation was indistinguishable under control or EFR3-knockdown conditions (Figure 3C). This observation suggests that the insulin-stimulated translocation of GLUT4 to the PM is not significantly impaired in EFR3-depleted cells, but the inability of GLUT4 to undergo insulin-stimulated dispersal underscores the impairment in insulin-stimulated glucose transport (Figure 2).

### Summary

The idea of localisation controlling the function and activity of PM resident proteins is well-established (Eisenberg et al., 2021; Kusumi et al., 2005, 2010, 2011; Trimble and Grinstein, 2015). Our work provides a link between EFR3 (and PI4K-IIIα) and insulin-stimulated glucose transport, establishing EFR3 as a key locus of insulin action. Analysis of GLUT4 distribution in the PM demonstrates that the insulin-stimulated dispersal of GLUT4 in the PM is dependent upon EFR3. A challenge for the future is to understand how this relates to GLUT4 dynamics in the PM, and how the insulin signalling machinery integrates with the EFR3 scaffold.

## Materials and methods

### Antibodies

Anti-GLUT4 and anti-Syntaxin 4 were from Synaptic Systems (Germany; #235003 and 110042, respectively). Anti-GAPDH was from ThermoFisher (Renfrew, UK; #AM4300). Anti-EFR3 was from Sigma (Dorset UK; #HPA023092) and anti-PI4K-IIIα was from AbCam (Cambridge, UK; #111565). Anti-HA was from Covance (Suffolk, UK; mouse monoclonal MMS101P) or from Roche (Wellwyn Garden City, UK; rat monoclonal 11867 423001). The Alexa Fluor 647-conjugated anti-HA antibody for dSTORM experiments was from Invitrogen (Renfrew, UK; mouse monoclonal 26183-A647). Secondary antibodies were from LICOR Biosciences (Cambridge UK; donkey anti-rabbit: 925 68023; goat anti-mouse 925 68070; goat anti-mouse 926 32210 and goat anti-rat [for FACS] A-21247).

### Plasmids

EFR3A- and EFR3B tagged with mCherry, and the corresponding tomato-tagged C(6-9)S mutant were provided by Pietro De Camilli (Yale University) and are described in (Chung et al., 2015; Nakatsu et al., 2012b).

### Growth and transfection and transport assays of 3T3-L1 adipocytes

3T3-L1 murine adipocytes were purchased from ATCC (via LGC Standards, USA) and grown and differentiated as outlined (Roccisana et al., 2013). Stable lines of 3T3-L1 fibroblasts expressing HA-GLUT4-GFP had previously been generated in the lab (Morris et al., 2020). Cells were incubated in a 10% CO_2_ humid atmosphere incubator at 37°C.

For electroporation, cells were used at day 6 post-differentiation. Cells were washed in PBS before detaching using 0.05% (w/v) trypsin: 2mg/ml collagenase. Once detached, cells were washed and transferred to 0.2cm BioRad Gene Pulser^®^ electroporation cuvette containing 3nmol Silencer^®^ select pre-designed siRNA and electroporated using settings of 0.18kV and 975μF. Cells were then plated and assayed 72 h after electroporation. siRNA against EFR3 was siRNA ID s94606 (ThermoFisher) and PI4K-IIIα siRNA ID s104706 (ThermoFisher).

2-deoxy-**d**-glucose (2DG) uptake was assayed as in (Roccisana et al., 2013). Non-specific association of radioactivity with the cells were quantified by performing parallel assays in the presence of 10 μM cytochalasin B (Bloch, 1973).

### Subcellular fraction and immunoblotting

10cm plates of 3T3-L1 adipocytes between day 8 and 10 were serum starved using serum free DMEM for 2 hours and incubated with or without insulin as in the figure legends. Plates were washed using sterile HES buffer, scraped and homogenised using a Dounce homogeniser and subcellular fraction performed as described (Roccisana et al., 2013). This procedure generates a fraction enriched in the PM (PM), high-density membranes (HDM), low density membranes (LDM) and soluble protein fraction (Piper et al., 1991; Simpson et al., 1983). Insulin results in a redistribution of GLUT4 from the LDM to the PM fraction. SDS-PAGE and immunoblotting for proteins within these fractions was performed as outlined (Roccisana et al., 2013).

### Semi-quantitative reverse transcriptase PCR

mRNA was extracted from the cells using QiaGEN mRNA easy kit as per the manufacturer’s instructions and mRNA quantified using a Nanodrop 1000. cDNA was prepared from the mRNA using Applied Biosystems reagents for RT-PCR, and then used in an Applied Biosystems Power SYBR-green PCR master mix.

### Cell preparation for dSTORM

Prior to measurement, 3T3-L1 adipocytes expressing HA-GLUT4-GFP were serum starved for 2 h. Cells were then stimulated with 100 nM insulin for 20 min or left untreated. Subsequently, cells were fixed with 4% paraformaldehyde (PFA) in PBS overnight at 4°C. The samples were quenched with 50 mM NH_4_Cl in PBS for 10 min at room temperature, washed with PBS then incubated in blocking solution (2% BSA with 5% goat serum in PBS) for 30 min. Afterwards cells were incubated with a conjugated anti-HA antibody coupled to Alexa Fluor 647 at a concentration of 8 μg/mL in blocking solution for 1 hour in the dark. Samples were washed with PBS for 10 minutes 3 times on an orbital shaker.

### dSTORM image acquisition and reconstruction

The dSTORM image sequences were acquired on an Olympus IX-81 microscope equipped with Olympus Cell^R acquisition software, an ImageEM EM-CCD 512 × 512 camera (Hamamatsu UK) and an Olympus × 150 UAPO oil lens with a numerical aperture of 1.45 and a resulting pixel size of 106 nm. Cold dSTORM imaging buffer containing 50mM mercaptoethylamine (MEA) in PBS (pH=8) was pipetted into cavity slides and coverslips were sealed onto slides using dental paste to avoid oxygen entry. After mounting of samples image sequences of 10,000 frames were acquired in total internal reflection fluorescence (TIRF) configuration using 647 nm laser light at 100% power (150 mW). Images were recorded on an Andor iXon 897 EMCCD camera using a centered 256 × 256 pixel region at 30 ms per frame for 10,000 frames and an electron multiplier gain of 200 and pre-amplifier gain profile 3. The dSTORM data were processed using the freely available Image J plugin ThunderSTORM (Ovesný et al., 2014). The image reconstruction parameters chosen are lined out in the following: pre-detection wavelet filter (B-spline, scale 2, order 3), initial detection by local maximum with 8-connected neighbourhoods (radius 1, threshold at two standard deviations of the F1 wavelet), and sub-pixel localisation by integrated Gaussian point-spread function (PSF) and maximum likelihood estimator with a fitting radius of 3 pixels. The first pass detected localisations were filtered according to the following criteria: an intensity range of 500 - 5000 photons, a sigma range of 25 - 250, and a localisation uncertainty of less than 25 nm.

Subsequently, the filtered data set was corrected for sample drift using cross-correlation of images from 5 bins at a magnification of 5. Repeated localisations, such as can occur from dye re-blinking, were reduced by merging points which re-appeared within 20 nm and 1 frame of each other.

### HDBSCAN analysis

The density-based spatial clustering of applications with noise (*DBSCAN*) has become one of the most common data clustering algorithms since its development (Ester et al., 1996). Density-based clustering defines clusters as areas of higher density compared to the remainder of the data points. Hierarchical *DBSCAN* (*HDBSCAN*) uses an unsupervised learning algorithm to find clusters of varying densities (Campello et al., 2013). HDBSCAN identifies regions of the data that are denser than the surrounding space and considers cluster hierarchy which is shaped by multivariate modes of the underlying distribution (Campello et al., 2013, 2015). ROI of 4μm x 4μm were selected in ImageJ for each cell and GLUT4 molecule coordinates were clustered using *HDBSCAN* parameters of *min_cluster_size=5* and *min-samples=30* to provide clear visualisation of the dataset. The *HDBSCAN* package (v0.8) was implemented on Python (v3.6) was downloaded from Github (https://github.com/scikit-learn-contrib/hdbscan) (McInnes et al., 2017).

### Spatial statistics analysis in R

Spatstat is a package for analysing spatial point pattern data featuring a generic algorithm for fitting point process models to point pattern data (Baddeley and Turner, 2005). Analysis of spatial statistics was carried out using the *spatstat* package (v1.64) for R (v3.2). GLUT4 clustering does not show uniform point density and statistics of point pattern correlation with correction for point pattern inhomogeneity and edge correction was applied. Cell outlines were chosen as ROI for the analysis using ImageJ. dSTORM localisations only within cells were further analysed. Ripley’s K function defines and measures covariance in a point process with clustered patterns showing positive covariance, independently placed points showing zero variance and dispersed points showing negative covariance (Baddeley et al., 2016).

The K function of a point process shows the expected number of neighbours of a point of a typical stationary point process **X** found at ***u*** that is within a distance ≤ ***r***:

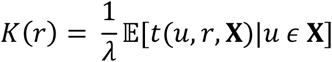

The empirical Ripley’s K function is obtained by measuring all the pairwise distances of points within one cell then standardised for the density of points.

The variance stabilised L function transforms K(r) into a straight line and is defined as:

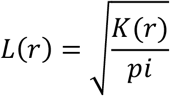

The empirical L(r) was calculated for each cell and pooled. For visualisation purposes, L(r) was centered, which is shown in the relevant figures. The experiment was carried out independently 4 times for a total of N=10 cells per group. Empirical estimates of the L function from each were pooled together to provide a weighted mean and the 83% (or 1.37σ) confidence interval for each group. Statistical differences between group means were assessed by overlap of 83% confidence intervals. In statistics checking for overlap of 83% confidence levels has been frequently reported as a method to assess whether two means are significantly different from each other or not at the α = 0.05 level (Austin and Hux, 2002; Goldstein and Healy, 1995; Payton et al., 2003).

### Localization density of GLUT4 molecules

Localization density of GLUT4 molecules in the PM was assessed using the Image J plugin LocFileVisualizer_v1.1 as described (van de Linde, 2019).

### Mice and preparation of heart and muscle samples

Mice used in this study were housed in single-sex groups of 5-6 mice per cage at the Central Research Facility at the University of Glasgow and kept on 12 h cycles of light and dark and at ambient temperature. Mice were purchased from Harlan Laboratories (Oxon, UK). All animal experiments were performed in accordance with the United Kingdom Home Office Legislation under the Animals (Scientific Procedure) Act 1986 (project licenses 60/4224 and 70/8572 which were approved by the Glasgow University Animal Welfare and Ethical Review Board) and guidelines from Directive 2010/63/EU of the European Parliament on the protection of animals used for scientific purposes.

Mice were randomly divided into two groups and fed either a normal diet or a high fat diet for 12 weeks starting at 8 weeks of age. The high-fat diet (Western RD) was purchased from SDS (SDS diets, U.K) and contained: fat 21.4%, protein 17.5% and carbohydrate 50% (42% kcal fat). Body weight was monitored weekly. At the end of 12 weeks mice were fasted for 16 h before a glucose tolerance test (Mancini et al., 2017) and were then euthanised by a rising concentration of CO_2_. Tissues were snap frozen in liquid nitrogen prior to preparation of homogenates as described (Mancini et al., 2017). At the end of the 12-week period of feeding there was a significant increase in body weight in WT mice on HFD compared to mice fed normal diet (see (Almabrouk et al., 2018) for details). Fasting blood glucose and plasma insulin measured at the end of the study period both showed a tendency toward an increase in the HFD group, as did incremental area under the curve following the glucose tolerance test (Almabrouk et al., 2018).

## Acknowledgements

This work was supported by the British Heart Foundation (Studentship to AMK: FS/14/61/31284), Diabetes Research and Wellness Foundation (grant SCA/PP/12/20 to GWG and NJB), (Diabetes UK (grant 15/000526 and 13/0004696 to GWG and NJB), and a Lord Kelvin Adam Smith studentship (to SM). NG and MFAC acknowledge ERC funding through FAKIR 648892 Consolidator Award. We thank Dr Anna White (University of Glasgow) for tissue samples, Pietro De Camilli (Yale University) for generous access to plasmid constructs, Leandro Lemgruber Soares (University of Glasgow) for help with dSTORM, and Sebastian Van De Linde (University of Strathclyde) and Dylan Owen (Kings College London) for helpful discussions. There are no conflicts of interest to disclose.

